# Nonlinear effect of light intensity on normal axial development of rhesus monkeys

**DOI:** 10.1101/2022.06.28.497947

**Authors:** Ying-Zhou Hu, Hua Yang, Jing Wu, Hao Li, Long-Bao Lv, Zhu Zhu, Lu-Yao Zhou, Yu-Hua Zhang, Fang-Fang Yan, Shu-Han Fan, Cheng-Yu Li, Shu-Xiao Wang, Jian-Ping Zhao, Qiang Qi, Chang-Bing Huang, Xin-Tian Hu

## Abstract

**Purpose:** To investigate the effects of different indoor lighting intensity (500 lx, 750 lx and 1,000 lx) on normal ocular axial length growth by using juvenal rhesus monkeys.

**Methods:** Twenty-four juvenile monkeys were exposed continuously to normal intensity light (NIL, 500 lx, n=16), medium intensity light (MIL, 750 lx, n=8) and high intensity light (HIL, 1 000 lx, n=8), with a same CCT value (about 3000 K) for 246 days. Axial length, anterior chamber depth, corneal curvature radius were measured at about a monthly interval.

**Results:** After 246 days of light exposure, the growth of axial length of the MIL group (750 lx) were 0.151 ± 0.081 mm and 0.139 ± 0.070 mm in the right and left eyes, respectively, and significantly larger in comparison with the NIL group (500lx, OD: 0.068 ± 0.055 mm, OS: 0.074 ± 0.057 mm) and the HIL group (1000lx,OD:0.063 ± 0.093 mm, OS: 0.084 ± 0.052 mm) monkeys. **This effect was stable and robust during the whole experimental period.**

**Conclusion:** The effects of different intensity lighting on normal ocular axial development was not linear as most people currently think. We must be cautious when it comes to elevate light intensity in classrooms. Whether this conclusion is correct under lights of other CCT value needs further study.

## 1. Introduction

In recent decades, the incidence of myopia has increased dramatically, especially among children in East Asia (Pan et al., 2012, Holden et al., 2016, Grzybowski et al., 2020). The incidence of juvenile myopia in China, South Korea and Japan is 52.7% (Mu et al., 2022), 73 % (Rim et al., 2016) and 76.5 % - 94.9 % (Yotsukura et al., 2019), respectively, ranking among the top in the world (Grzybowski et al., 2020). This grim situation has caused great international concern worldwide.

Recent studies from multiple laboratories have shown that outdoor activities can slow down the development of myopia in children (Jones et al., 2007, Rose et al., 2008, Dirani et al., 2009, Deng et al., 2010, Guggenheim et al., 2012, French et al., 2013). But the outdoor and indoor environments are quite different, and therefore, it is not clear what factors slow down the development of myopia. One possible reason is the huge different light intensity between indoor and outdoor. The illuminance of outdoor could exceeds 100,000 lx, hundreds of times higher than that of indoor (typically no more than 500 lx). To test this hypothesis, researchers conducted several experiments on a variety of experimental animals. When exposed to high intensity light (between 10,000 to25,000 lx, equivalent to the light intensity level under tree shade on a sunny day), the elongation of ocular axis and myopic shift of chickens, tree shrews and macaques significantly slowed down than exposed to normal indoor intensity light (100-500 lx) (Cohen et al., 2011, Siegwart et al., 2012, Smith et al., 2012). In other words, in the three species, compared with normal indoor light, exposure to high intensity outdoor light would slow the progression of myopia. Studies on children have also shown that myopic incidence and axial length are significantly lower and shorter in children exposed to higher illuminance lights than in groups of children exposed to lower illuminance lights (Hua et al., 2015, Zhou et al., 2017, Wu et al., 2018). Therefore, it is generally accepted that high level illuminance lighting may be one of the reasons why increasing outdoor activities time can reduced the incidence of myopia (Norton and Siegwart Jr, 2013, Chen et al., 2017, Mohanty et al., 2021).

In general lighting, in addition to the light intensity, correlated color temperature (CCT) and color rendering index (CRI) are other two important parameters that determine lighting quality. The relationship between CCT and the development of myopia has not been reported till recently. Axial growth rate is an important and reliable indicator of the development and progression of myopia (Rucker, 2019). Since rhesus monkeys and humans are very close in evolution, the structure and function of the eyes of rhesus monkeys and humans are very similar, and the possibility of effective extrapolation of experimental results from monkeys to humans is highly guarantied (Morgan et al., 2014, Schaeffel and Feldkaemper, 2015). Recently, we (2022) had reported the first research on the effects of commercially available CCT on the development of ocular axial length in juvenile rhesus monkeys. The results demonstrate that, compared with high CCT light (4000 K and 5000 K), low CCT light (2700 and 3000 K) exposure can significantly slow down the elongation of axial length in juvenile rhesus monkeys, and this effect is robust and stable throughout the whole experiment (Hu et al., 2022). Thus, this interdisciplinary study has filled an important gap in the research of general lighting and ocular development and proposed a new method of school myopia control and prevention by maintaining the CCT of indoor lighting around 3000K.

The above results were obtained when the light intensity was set at 500 lx. In current study, in order to further enhance the effect of myopia prevention of general lighting, the color temperature was set at 3000K and the relationship between light intensity and ocular axis development was systematically explore using juvenal monkeys. In the past studies on the relationship between the axial length development and light intensity, to mimic outdoor light, the intensities were usually exceeded 10,000 lx, which is very hard to apply in daily indoor lighting. The purpose of this study was to find optimal light intensity within the range of daily lighting which is usually below 1000 lx, so we varied the light intensity of this study at: 500, 750, and 1000 lx. Through strict quantitative macaque experiments, we had found an interesting results on the relationship of ocular axis development and.

After a rigorous juvenal monkey experiment, we made unexpected important discovery on the relationship between the ocular axis development and different light different intensity exposure levels within 500-1000 lx range of a commercially available 3000 K lighting.

## 2. Methods

### 2.1 Animals and welfare

Thirty-two healthy young rhesus macaques (Macaca mulatta) (16 males, 16 females) were purchased from the Kunming Primate Research Center, Chinese Academy of Sciences (Yunnan, China). The monkeys were 3.18–3.45 years old (3.33±0.07 years; mean ± SD), which are comparable to Grade 4-Grade 6 primary school children (9–12 years). All rearing and experimental procedures were reviewed and approved by the Institutional Animal Care and Use Committee (IACUC) of the Kunming Primate Research Center (Approval Number: IACUC20029) and were in strict compliance with the National Care and Use of Animals Guide approved by the National Animal Research Authority (China) and the National Institutes of Health Guide for the Care and Use of Laboratory Animals (USA).

### 2.2 Intervention protocol

Measurements of ocular axial length, corneal curvature radius, anterior chamber depth, and spherical equivalent (SE) refractive errors were performed on the monkeys before the experiment and then regularly after the experiment started. Ocular measurements were carried out following standard protocols used in previous studies (Ivers et al., 2011, Qiao-Grider et al., 2007). All data were collected between 1200 h and 1630 h on testing days.

The monkeys were exposed to commercially available LED light (CCT, ~ 2800K) at three different levels of intensity. They were: Group 1 (the normal intensity light (NIL) group (n=16), with an intensity of ~ 500 lx); Group 2 (the medium intensity light (MIL) group (n=8), with an intensity of ~ 750 lx) and Group 3 (the high intensity light (HIL) group (n=8), with an intensity of ~ 1,000 lx). The mean axial lengths at the beginning of the experiments were comparable among groups (Table 1, F2, 29=0.132, P=0.877). Eight single monkey cages arranged in two columns were housed in each experimental room. All animals were transferred into the single cages (0.8 × 0.8 × 0.8 m) in the experimental rooms on the same day, one in each cage. During the experiment, illumination at the center of each cage was maintained at ~500 lx, ~750 lx and ~1,000 lx for the corresponding group by careful measurements and adjustments at regular intervals (29–34 days). The daily light cycle was 12 h light/12 h dark, with lights on from 0700 h to 1900 h. The monkeys were only exposed to artificial light after transfer to the lighting room. All animals were supplied with food and water ad libitum and were inspected daily by experienced veterinarians.

Both eyes of monkey were not restricted during light exposure. They developed naturally under artificial lighting to mimic the developmental process of the ocular axis in children under indoor activity conditions.

### 2.3 Data collection

Measurements of ocular axial length, corneal curvature radius, anterior chamber depth, and refractive status (spherical equivalent, SE) were performed on the monkeys before the experiment and then regularly after the experiment started. Each monkey was anesthetized with ketamine (intramuscular administration, 20–25 mg/kg) and acepromazine maleate (intramuscular administration, 0.15–0.20 mg/kg). Ocular axial length, corneal curvature radius, and anterior depth were then measured using an optical biometer (IOLMaster, Carl Zeiss Meditec, Germany). To prepare for refractive status collection, cycloplegia and mydriasis were induced by tropicamide phenylephrine eye drops (Mydrin-p, Santen, Osaka, Japan) instilled 25 and 20 mins before the measurements. Refractive status examination was performed using a computerized autorefractor (KR-800, TOPCON Medical Systems Inc., Japan). Due to the similarity between monkey and human eyes, refractive status of monkey eyes was measured by optical biometry and computerized autorefractometry,which has been proven feasible and effective in monkeys (Hung et al., 2012, Ostrin et al., 2012, Lin et al., 2019).

### 2.4 Data analysis

Data from both eyes of each monkey were analyzed and controlled as within-subject effect. The trends of ocular parameters changes over the experimental time were analyzed and compared using generalized linear mixed model (GLMM) follow by Bonferroni correction for post hoc tests, which is commonly used in the field (Lin and Tsai, 2016, Lee et al., 2020). At the end of the experiments, the last set of ocular data were analyzed by repeated measures one-way analysis of variance (repeated measures ANOVA), followed by LSD post hoc tests, and the two eyes’ data were controlled as within subject effect(Armstrong, 2013). All statistical analyses were performed using SPSS Statistics v27.0 (IBM Corp., Armonk, NY, USA).

## 3. Results

The data demonstrated that LED lights with different intensity had different effects on ocular axial growth in the juvenile monkeys (repeated measures ANOVA: *F*_2, 29_ = 4.135, *P* = 0.026, Figure 1A). After 246 days of light exposure at the end of the experiment, the changes of axial length of the MIL group(750lx) were 0.151 ± 0.081 mm and 0.139 ± 0.070 mm in the right and left eyes, respectively, and were significantly larger in comparison to the NIL group (500lx,OD:0.068 ± 0.055 mm, OS: 0.074 ± 0.057 mm) and the HIL group (1000lx,OD:0.063 ± 0.093 mm, OS: 0.084 ± 0.052 mm) monkeys (ANOVA fellow by LSD, *P* = 0.011,*P* = 0.028, respectively). There were no significant differences of changes of axial length between the NIL group and HIL group (ANOVA fellow by LSD, P = 0.945) (Figure 1 A).

**Figure 1.**
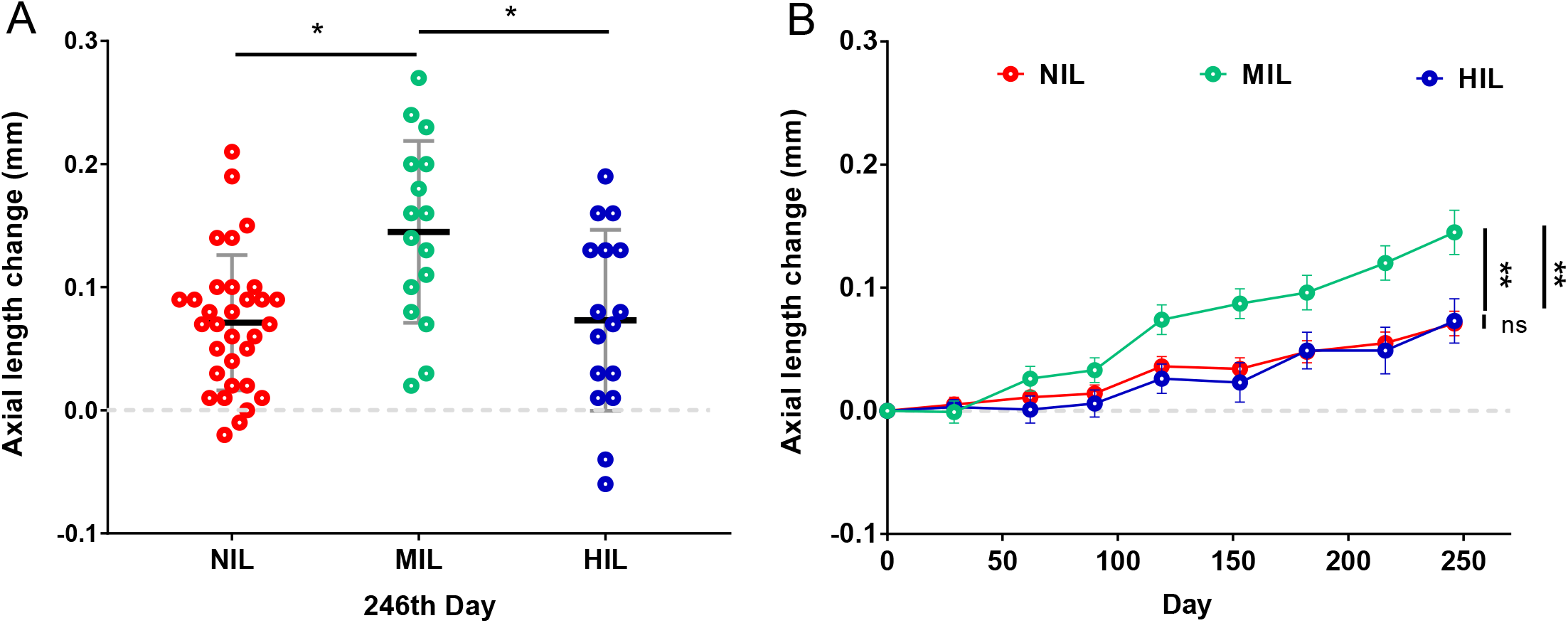
Ocular axial length elongation of juvenal monkeys under different intensities of LED lighting after 246 days of exposure. A: The axial length changes of MIL group (750 lx, green circle) was significantly higher than NIL group (500 lx, red circle) and HIL group (1000 lx, blue circle). B: Increased ocular axial length over time under three different light intensity. Ocular axial growth rate of monkeys’ exposure to medium intensity light (750 lx) was higher than those of monkeys exposure to normal intensity light and high intensity light. Differences in growth rates of Ocular axial length between NIL group and HIL group were not significant (ns). *: *P*<0.05; ** *P*<0.01. Data are mean ± SD in A and mean ± SEM in B.

Significant differences in the rates of increasing trend of ocular axial length during the whole experimental period were also observed among the three intensity groups (Figure 1B). Using generalized linear mixed models (GLMM), we found that intensity had a significant effect on the trend of ocular axial growth during the whole lighting period (GLMM: *F*_2, 509_ = 13.718, *P* < 0.001; Fig. 1 B). Paired comparative analysis of the GLMM further demonstrated that the ocular axial growth rate of monkeys under medium intensity light was higher than those of monkeys’ exposure to normal intensity light (t = 4.353, p < 0.001; Bonferroni) and high intensity light (Bonferroni: t = 4.892, p < 0.001). In addition, there was no significant difference in ocular axial growth rate between NIL and HIL group (t = 1.295, p =0.196).

## 4. Discussion

In this study, we examined the effects of three light intensities (500 lx, 750 lx, 1000 lx) of commercially available LED with a CCT of 3000K on normal ocular growth of juvenile rhesus monkeys. After an indoor light exposure of 246 days, we found that the effects of different intensity lighting on normal ocular axial development was not linear. The growth rate of axial length of the yang monkeys were faster after exposed to the medium intensity light (MIL, 750 lx) than to the normal intensity light (NIL, 500 lx) and high intensity light (HIL, 1000 lx) (Fig. 1 A, B). The change of axial length of MIL group was almost twice as much than NIL and HIL group (Fig. 1 A). And the effects were robust and stable during the whole experimental period (Fig. 1 B)

This is an interesting and unexpected result. Why has this phenomenon not been reported in previous studies on the relationship between light intensity and the development of ocular axis? First, there were only a few studies has been done on light intensity and normal ocular growth. Most researchers used induced myopia animals models to study the effects of light intensity on ocular length growth (Norton, 2016), which are different from ours both in purpose and methodology. We searched on PubMed with “light intensity”, “illuminance” and “myopia”, and after excluded studies employed monochromatic light and induced myopia animal models, only 4 articles came out as studies on ocular development under different artificial lighting intensities. Among the 4 articles, one reports the effect of elevated light level on ocular development in primary school children (Hua et al., 2015), and the three other articles report the effect of different intensity lighting on ocular growth in experiment animals (Cohen et al., 2011, Cohen et al., 2012, She et al., 2020). The intensity levels in all these studies were not overlap with ours. Cohen et al. (2011, 2012) exposed chicks to 50 lx, 500 lx and 10,000 lx ambient lighting. The axial length of chicks under 10,000 lx and 500 lx light was significantly shorter than chicks’ under 50 lx light (Cohen et al., 2011, Cohen et al., 2012). They did not observe the non-linear effect of light intensity on ocular development could due to the huge differences among the exponential light intensity levels:10,000 lx is 20 folds and 200 folds to 500 lx and 50 lx. She et al. (2020) exposed infant monkeys to two intensity level (55 lx and 504 lx) lights and found no significant differences of axial elongation between the two groups (She et al., 2020). This result is at odds with Cohen et al (2011, 2012) from chicks’. We speculate that the difference in species is an important reason for this inconsistence. Hua et al. (2015) employed two intensity level (74 lx and 558 lx) in their study and found that pupils who studied under 558 lx had a significantly smaller increase in ocular axis than those who under 74 lx. However, their results are not comparable with ours for when only two intensity light levels are employed, it is impossible to observe a non-linear effect of light intensity on axial elongation.

Although this result was unexpected, it is in line with some recent studies. It is generally accepted that light exposure stimulates the retina to secrete dopamine-a stop signal of eye growth (karouta and Ashby, 2015, Landis et al., 2021) and that there is a dose-response relationship with light intensity (Brainard and Morgan, 1987). Further studies have pointed out that this relationship is not linear. Intermediate illuminance level light inhibits dopamine secretion by the mouse retina, whereas low - and high illuminance light induce higher levels of dopamine secretion (P é rez fern á ndez et al., 2019). Landis et al. (2021) found that lens induce myopia mice developed significantly less severe myopic refractive shifts when exposed to scotopic or photopic lighting than mice exposed to mesopic lighting. Although the species and specific light intensity are different, these recent research results are consistent with our current findings.

Our results indicate that there are flaws in the current commonly held assumption that, in the daily illumination range, the stronger the light intensity, the slower the axial growth of the eye. When it comes with the increase of light intensity, extra caution should be taken. Should it be just 500 or 1000 lx? Not necessarily. One limitation of our experiment is that the relationship between light intensity and ocular axial development was only tested at a color temperature of 3000 K. Whether the conclusions are correct under other chromatic temperature conditions awaits further experimental investigation.

## Declarations

## Acknowledgements

This work was supported in part by the Key-Area Research and Development Program of Guangdong Province (2019B030335001), Science and Technology Service Network Initiative (STS) Project of Chinese Academy of Sciences (E02E1801), National Key R&D Program of China (2018YFA0801403), the Scientific Instrument Developing Project of the Chinese Academy of Sciences (CAS) (022006), the Strategic Priority Research Program of the Chinese Academy of Sciences (XDB32060200), the National Natural Science Foundation of China (81941014, 81771387, 31800901, 31960178), CAS “Light of West China” program, the Applied Basic Research Programs of Science and Technology Commission Foundation of Yunnan Province (202001AT070130), and National Research Facility for Phenotypic and Genetic Analysis of Model Animals, Kunming Institute of Zoology, Chinese Academy of Sciences.

We would like to thank Mr. Zheng-Fei Hu for the daily care of the monkeys, and Mr. Shun-Long Wang for his hard work on data collection.

## Ethical approval

The experiments were performed as per approval of the Institutional Animal Care and Use Committee (IACUC) of the Kunming Primate Research Center. Approval Number: IACUC20029

## Availability of data and materials

The datasets used and/or analysed in the present study can be obtained from the corresponding author on request.

## Authors’ contributions

Investigation: Y.Z.H., H.Y., H.L., L.B.L., J.W., Z.Z., and Y.H.Z. performed the experiments, Y.Z.H., F.F.Y., S.H.F., L.Y.Z. analyzed the data, X.T.H, H.Y., Q.Q., C.B.H., J.P.Z., S.X.W., C.Y.L. designed the project. Y.Z.H, X.T.H, and H.Y. wrote the manuscript, C.B.H., S.X.W., J.P.Z., H.L., and X.T.H. revised the manuscript. All authors read and approved the final version of the manuscript.

## References

Armstrong R A 2013. Statistical guidelines for the analysis of data obtained from one or both eyes. Ophthalmic and Physiological Optics [J], 33: 7–14.

Chen S, Zhi Z, Ruan Q, et al. 2017. Bright light suppresses form-deprivation myopia development with activation of dopamine D1 receptor signaling in the ON pathway in retina. Investigative ophthalmology & visual science [J], 58: 2306–2316.

Cohen Y, Belkin M, Yehezkel O, et al. 2011. Dependency between light intensity and refractive development under light–dark cycles. Experimental eye research [J], 92: 40–46.

Cohen Y, Peleg E, Belkin M, et al. 2012. Ambient illuminance, retinal dopamine release and refractive development in chicks. Experimental eye research [J], 103: 33–40.

Deng L, Gwiazda J, Thorn F 2010. Children’s refractions and visual activities in the school year and summer. Optometry and Vision Science [J], 87: 406.

Dirani M, Tong L, Gazzard G, et al. 2009. Outdoor activity and myopia in Singapore teenage children. British Journal of Ophthalmology [J], 93: 997–1000.

French A N, Ashby R S, Morgan I G, et al. 2013. Time outdoors and the prevention of myopia. Experimental eye research [J], 114: 58–68.

Grzybowski A, Kanclerz P, Tsubota K, et al. 2020. A review on the epidemiology of myopia in school children worldwide. BMC ophthalmology [J], 20: 1–11.

Guggenheim J A, Northstone K, Mcmahon G, et al. 2012. Time outdoors and physical activity as predictors of incident myopia in childhood: a prospective cohort study. Investigative ophthalmology & visual science [J], 53: 2856–2865.

Holden B A, Fricke T R, Wilson D A, et al. 2016. Global prevalence of myopia and high myopia and temporal trends from 2000 through 2050. Ophthalmology [J], 123: 1036–1042.

Hua W J, Jin J X, Wu X Y, et al. 2015. Elevated light levels in schools have a protective effect on myopia. Ophthalmic and Physiological Optics [J], 35: 252–262.

Hung L-F, Ramamirtham R, Wensveen J M, et al. 2012. Objective and subjective refractive error measurements in monkeys. Optometry and vision science: official publication of the American Academy of Optometry [J], 89: 168.

Ivers K M, Li C, Patel N, et al. 2011. Reproducibility of measuring lamina cribrosa pore geometry in human and nonhuman primates with in vivo adaptive optics imaging. Investigative ophthalmology & visual science [J], 52: 5473–5480.

Jones L A, Sinnott L T, Mutti D O, et al. 2007. Parental history of myopia, sports and outdoor activities, and future myopia. Investigative ophthalmology & visual science [J], 48: 3524–3532.

Lee M W, Lee S-E, Lim H-B, et al. 2020. Longitudinal changes in axial length in high myopia: a 4-year prospective study. British Journal of Ophthalmology [J], 104: 600–603.

Lin C-J, Tsai Y-Y 2016. Axial length, refraction, and retinal vascularization 1 year after ranibizumab or bevacizumab treatment for retinopathy of prematurity. Clinical ophthalmology (Auckland, NZ) [J], 10: 1323.

Lin R, Wu K-C, Jin G-H, et al. 2019. Screening of high myopia in non-human primates. Investigative ophthalmology & visual science [J], 60: 5870–5870.

Mohanty P, Brown D, Mazade R, et al. 2021. Rod pathway signaling has protective effects on myopia susceptibility in dim, but not bright light. Investigative ophthalmology & visual science [J], 62: 1395–1395.

Morgan I G, Rose K A, Ashby R S 2014. Animal models of experimental myopia: limitations and synergies with studies on human myopia. Pathologic myopia [J]: 39–58.

Mu J, Zhong H, Liu M, et al. 2022. Trends in Myopia Development Among Primary and Secondary School Students During the COVID-19 Pandemic: A Large-Scale Cross-Sectional Study. Frontiers in Public Health [J], 10.

Norton T T 2016. What Do Animal Studies Tell Us about the Mechanism of Myopia– Protection by Light? Optometry and vision science: official publication of the American Academy of Optometry [J], 93: 1049.

Norton T T, Siegwart JR J T 2013. Light levels, refractive development, and myopia–a speculative review. Experimental eye research [J], 114: 48–57.

Ostrin L A, Hung L-F, Smith III E L, et al. 2012. Lamina Cribrosa Geometry and Axial Length in Normal Rhesus Monkeys. Investigative ophthalmology & visual science [J], 53: 3469–3469.

Pan C, Ramamurthy D, Saw S 2012. Worldwide prevalence and risk factors for myopia. Ophthalmic and Physiological Optics [J], 32: 3–16.

Qiao-Grider Y, Hung L-F, Kee C-S, et al. 2007. A comparison of refractive development between two subspecies of infant rhesus monkeys (Macaca mulatta). Vision Research [J], 47: 1668–1681.

Rim T H, Kim S-H, Lim K H, et al. 2016. Refractive errors in Koreans: the Korea National Health and nutrition examination survey 2008-2012. Korean Journal of Ophthalmology [J], 30: 214–224.

Rose K A, Morgan I G, Ip J, et al. 2008. Outdoor activity reduces the prevalence of myopia in children. Ophthalmology [J], 115: 1279–1285.

Schaeffel F, Feldkaemper M 2015. Animal models in myopia research. Clinical and Experimental Optometry [J], 98: 507–517.

She Z, Hung L-F, Arumugam B, et al. 2020. Effects of low intensity ambient lighting on refractive development in infant rhesus monkeys (Macaca mulatta). Vision Research [J], 176: 48–59.

Siegwart J T, Ward A H, Norton T T 2012. Moderately Elevated Fluorescent Light Levels Slow Form Deprivation and Minus Lens-Induced Myopia Development in Tree Shrews. Investigative ophthalmology & visual science [J], 53: 3457–3457.

Smith E L, Hung L, Huang J 2012. Protective Effects of High Ambient Lighting on the Development of Form-Deprivation Myopia in Rhesus Monkeys. Investigative ophthalmology & visual science [J], 53: 421–428.

Wu P-C, Chen C-T, Lin K-K, et al. 2018. Myopia prevention and outdoor light intensity in a school-based cluster randomized trial. Ophthalmology [J], 125: 1239–1250.

Yotsukura E, Torii H, Inokuchi M, et al. 2019. Current prevalence of myopia and association of myopia with environmental factors among schoolchildren in Japan. JAMA ophthalmology [J], 137: 1233–1239.

Zhou Z, Chen T, Wang M, et al. 2017. Pilot study of a novel classroom designed to prevent myopia by increasing children’s exposure to outdoor light. PloS one [J], 12: e0181772.

